# Stoichiometry of two plant glycine decarboxylase complexes and comparison with a cyanobacterial glycine cleavage system

**DOI:** 10.1101/2020.03.16.993188

**Authors:** Maria Wittmiß, Stefan Mikkat, Martin Hagemann, Hermann Bauwe

**Affiliations:** Department of Plant Physiology, University of Rostock, Albert-Einstein-Straße 3, D-18059 Rostock, Germany; Core Facility Proteome Analysis, Rostock University Medical Center, Schilling-Allee 69, D-18057 Rostock, Germany

**Keywords:** glycine cleavage system, glycine decarboxylase complex, multienzyme metabolic complexes, one-carbon metabolism, photorespiration, recombinant enzymes, *Synechocystis*

## Abstract

The multienzyme glycine cleavage system (GCS) converts glycine and tetrahydrofolate to the one-carbon compound 5,10-methylenetetrahydrofolate, which is of vital importance for most if not all organisms. Photorespiring plant mitochondria contain very high levels of GCS proteins organised as a fragile glycine decarboxylase complex (GDC). The aim of this study is to provide mass spectrometry-based stoichiometric data for the plant leaf GDC and examine whether complex formation could be a general property of the GCS in photosynthesizing organisms. The molar ratios of the leaf GDC component proteins are 1L_2_-4P_2_-8T-26H and 1L_2_-4P_2_-8T-20H for pea and Arabidopsis, respectively, as determined by mass spectrometry. The minimum mass of the plant leaf GDC ranges from 1,550-1,650 kDa, which is larger than previously assumed. The Arabidopsis GDC contains four times more of the isoforms GCS-P1 and GCS-L1 in comparison with GCS-P2 and GCS-L2, respectively, whereas the H-isoproteins GCS-H1 and GCS-H3 are fully redundant as indicated by their about equal amounts. Isoform GCS-H2 is not present in leaf mitochondria. In the cyanobacterium *Synechocystis* sp. PCC 6803, GCS proteins are present at low concentration but above the complex formation threshold reported for pea leaf GDC. Indeed, formation of a cyanobacterial GDC from the individual recombinant GCS proteins *in vitro* could be demonstrated. Presence and metabolic significance of a *Synechocystis* GDC *in vivo* remain to be examined but could involve multimers of the GCS H-protein that dynamically crosslink the three GCS enzyme proteins, facilitating glycine metabolism by the formation of multienzyme metabolic complexes.

## INTRODUCTION

The oxidation of glycine is vitally important for plants and most other organisms because it provides one-carbon-units to a large number of biosynthetic pathways (Engel *et al.*, 2007, Kikuchi *et al.*, 2008). The biochemical process requires three enzymes acting sequentially to produce 5,10-methylenetetrahydrofolate from glycine and tetrahydrofolate (THF; Figure 1). The three enzymes are the pyridoxal 5’-phosphate (PLP)-dependent P-protein (glycine decarboxylase, EC 1.4.4.2), the THF-dependent T-protein (aminomethyltransferase, EC 2.1.2.10), and the NAD^+^-dependent L-protein (dihydrolipoamide dehydrogenase, EC 2.1.8.1.4). P-, T- and L-protein share a common substrate protein, the H-protein (hydrogen or aminomethyl carrier protein) carrying a covalently linked lipoyl cofactor. During the reaction cycle, the lipoyl moiety of H-protein occurs in three different forms: the dithiolane form is the oxidant during glycine decarboxylation by P-protein, the aminomethylated form links the P-to the T-protein, and a dithiol form that is generated by T-protein. The latter must be re-oxidised by L-protein before the next reaction cycle can begin. Collectively, the four proteins form the glycine cleavage system (GCS). The GCS reaction cycle is fully reversible and can produce glycine from 5,10-methylenetetrahydrofolate, CO_2_, NH_3_ and NADH (Kawasaki *et al.*, 1966, Freudenberg and Andreesen, 1989). Hence, the GCS is being used in synthetic biology as a key component of the ‘reductive glycine pathway’ of formate and CO_2_ assimilation in tailor-made microbial organisms for sustainable bioproduction (discussed in Bar-Even, 2016, Yishai *et al.*, 2018).

**Figure 1.**
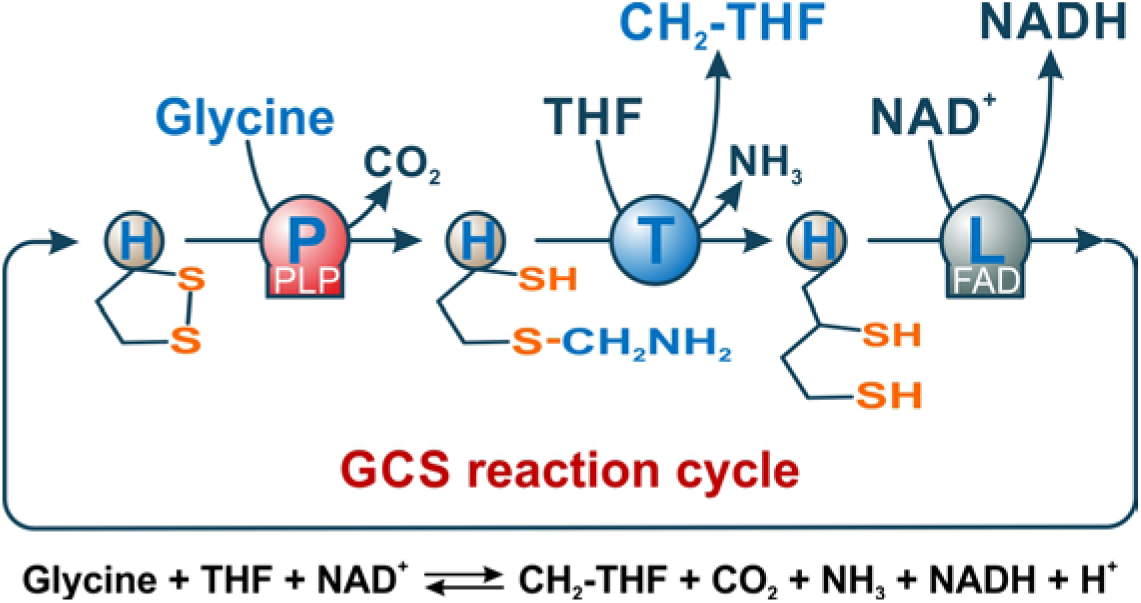
The GCS reaction cycle. Three closely cooperating enzymes, P-, T- and L-protein, oxidise glycine to form 5,10-methylene-THF, CO_2_ and NH_3_, reducing NAD^+^ to NADH. They produce and use three variants of the lipoyllysine arm of their shared substrate, H-protein (Kikuchi *et al.*, 2008). The dithiolane form serves as an oxidant and conveys the glycine’s methylene group to THF.

In eukaryotes, the GCS occurs only in the mitochondrion (Kisaki *et al.*, 1971, Motokawa and Kikuchi, 1971), closely cooperating with isoforms of SHMT located in different cellular compartments (for example, Bourguignon *et al.*, 1988) in order to provide one-carbon units and NAD(P)H for many biosynthetic pathways (for example, Mouillon *et al.*, 1999, Fan *et al.*, 2014). In photosynthesizing leaf cells, the GCS is mostly engaged in serine synthesis during photorespiration (reviewed in Bauwe *et al.*, 2010). The very high flux through this pathway requires unusally large amounts of GCS proteins in the mitochondrial matrix, about 130 mg ml^-1^ in pea leaf mitochondria (32% of total matrix protein mass), where they associate in the glycine decarboxylase complex (GDC, Neuburger *et al.*, 1986, Oliver *et al.*, 1990, Douce *et al.*, 2001). It is thought that formation of the GDC gives an activity boost to the GCS reaction cycle in order to match the needs of photosynthetic-photorespiratory metabolism. The latest hypothesis on the structure of the GDC speculates that about 30 H-protein molecules could form a central core to which one dimer of L-protein, two dimers of P-protein and nine monomers of T-protein are attached (Oliver and Raman, 1995). The underlying stoichiometry was determined by enzyme-linked immunosorbent assays (ELISA) protein quantification in the crude mitochondrial matrix extract in combination with activity measurements (Oliver *et al.*, 1990). By contrast to ‘classical’, stable multiprotein complexes, such as the 2-oxoacid dehydrogenase complexes or Rubisco, the GDC disaggregates easily at low concentration. Therefore, the GDC may be rather considered a multienzyme metabolic complex similar to those identified in glycolysis and the tricarboxylic acid cycle (reviewed in Schmitt and An, 2017). This view would correspond to the fact that the GCS is likewise operational at very low concentration *in vitro* and *in vivo*, for example in the mitochondria of heterotrophically cultured plant cells (about 0.18% of the mitochondrial proteome mass, Fuchs *et al.*, 2019) and animal cells as well as in prokaryotes. Information on the mitochondrial or cellular GCS concentration and hence the mode of GCS operation including a possible GDC formation in animals and prokaryotes is missing so far.

The objectives of this study are to examine the composition of plant GDCs by mass spectrometry technology and find out whether the potential to form a multiprotein complex is a general property of the GCS in photosynthesizing organisms. To this end, we investigated the stoichiometry and isoprotein composition of the *Arabidopsis thaliana* (Arabidopsis) leaf mesophyll GDC in comparison with that of *Pisum sativum* (pea) and examined whether and under which conditions recombinant GCS proteins of the cyanobacterium *Synechocystis* sp. PCC 6803 (*Synechocystis*) would be able to form a GDC.

## RESULTS

### Pea and Arabidopsis GDC differ from one another in their H-protein contents

First, by using a Hi3 shotgun proteomic approach, we confirmed the purity of the mitochondrial preparations and calculated a GDC content of ∼39 mole % (∼44 mass %) and an SHMT content of ∼13 mole % (∼14 mass%) in the matrix of Arabidopsis leaf mitochondria (Supplementary Table S 1), which is even higher than previous estimates for pea leaf mitochondria (∼30 mass %, Oliver *et al.*, 1990, Vauclare *et al.*, 1996). The mass spectrometry-based calculation of the GDC content of pea mitochondria was not possible, because a complete protein sequence database of pea is not available.

In order to quantify the molar ratios in which the leaf GDC proteins are present in pea and Arabidopsis mitochondria, we next designed an artificial protein comprising a concatenation of proteotypic tryptic peptides (QconCAT, Pratt *et al.*, 2006; Supplementary Figure S 1). After proteolytic digestion of the isotope-labeled QconCAT with trypsin, all expected peptides were identified (Supplementary Figure S 2) and showed a linear response with increasing concentration. Using total soluble mitochondrial matrix proteins spiked with the QconCAT, molar ratios of about 1L_2_-4P_2_-8T-26H and 1L_2_-4P_2_-8T-20H were calculated for pea and Arabidopsis, respectively (Figure 2). Comparable stoichiometries were obtained using isotope-labeled synthetic peptides, which however failed to quantify the P-protein, and label-free Hi3 quantification, which underestimated the amount of the H-proteins.

**Figure 2.**
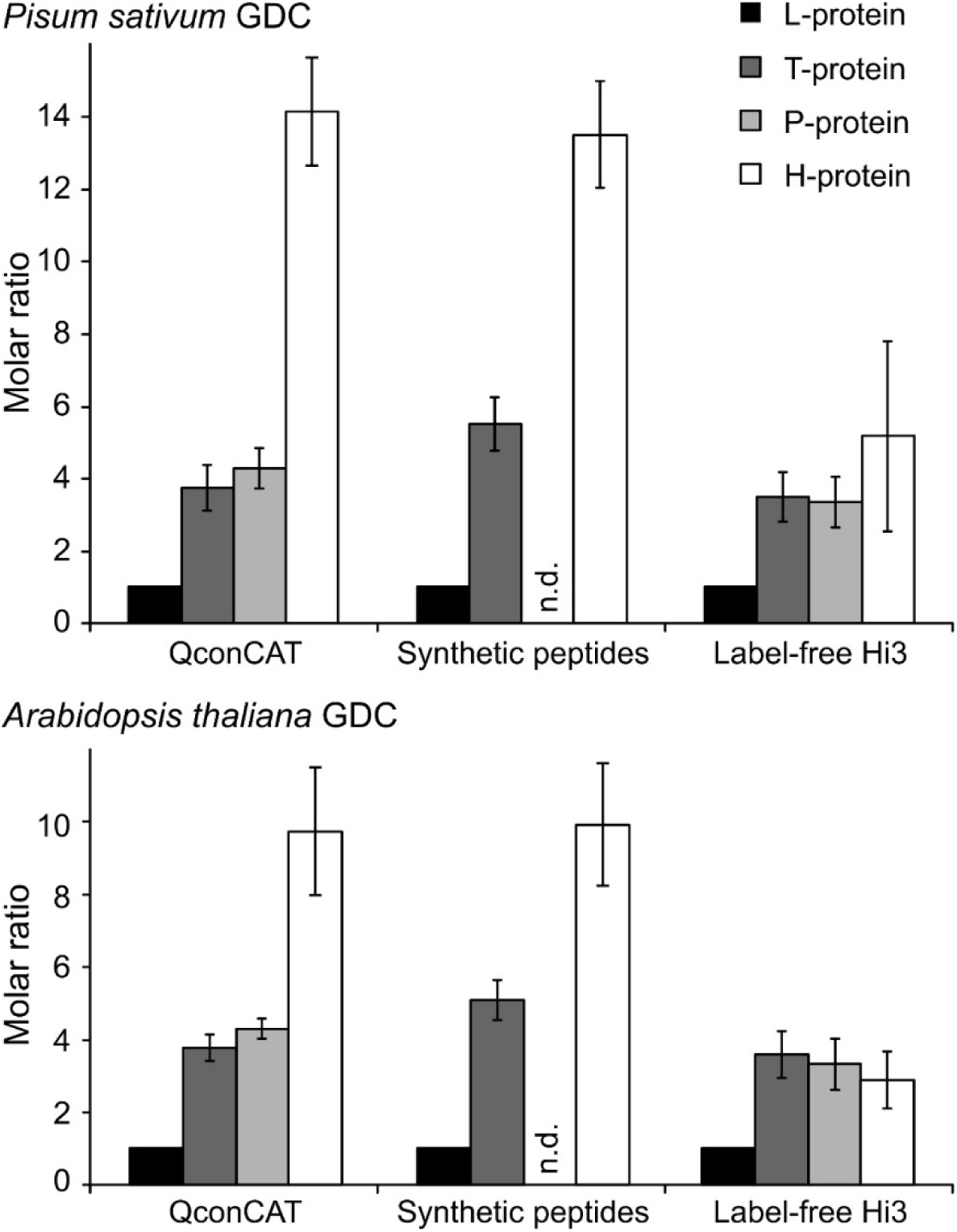
Stoichiometry of the GDC in pea and Arabidopsis leaf mitochondria. Molar ratios of the T-, P-, and H-proteins referred to L-protein (set to 1) are shown. Results from label-based quantification approaches using QconCAT and synthetic labeled peptides (SpikeTides) are compared to the label-free Hi3 method. No result is given for P-protein quantification with SpikeTides (n.d.), because of significant oxidation of the respective peptide preventing its correct quantification.

The P-protein, L-protein and H-protein are encoded by two (P and L) and three gene copies (H) each in the Arabidopsis genome (Bauwe and Kolukisaoglu, 2003), but the relative contribution of the respective isoforms to GCS activity and GDC formation is not known. Our QconCAT-based or label-free approaches were designed to determine the GDC protein isoform ratios in Arabidopsis leaf mesophyll mitochondria (Figure 3). The H-protein isoforms GCS-H1 (At2g35370) and GCS-H3 (At1g32470) are present in similar amounts, whereas the H-protein isoform GCS-H2 (At2g35120) was virtually absent. By contrast, P-protein and L-protein are mostly, each to about 80%, represented by the isoforms GCS-P1 (At4g33010) and mtLPD1 (At1g48030). The far most abundant mitochondrial SHMT is SHMT1 (At4g37930). The very low level of SHMT2 (At5g26780), which normally does not occur in mitochondria of photosynthesizing leaf cells but dominates in those of heterotrophic cells (Engel *et al.*, 2011, Fuchs *et al.*, 2019), indicates that preparations of leaf mesophyll mitochondria are inevitably contaminated by small amounts of mitochondria originating from other leaf tissues, such as the vasculature.

**Figure 3.**
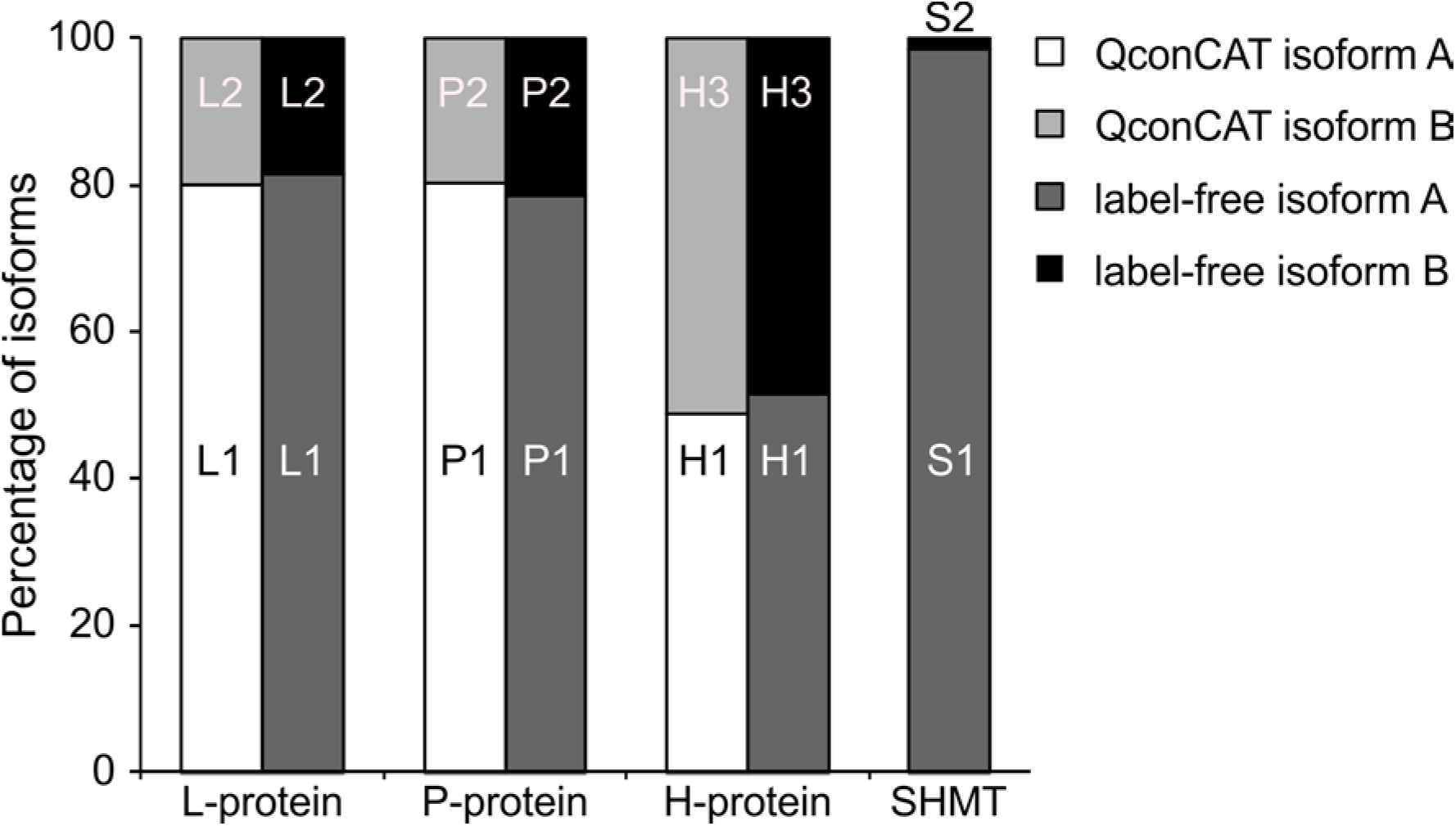
Isoproteins contributing to the GDC and to SHMT in Arabidopsis *leaf mitochondria.* Molar percentages of the GDC isoproteins were calculated by a label-free approach using all pairs of highly similar peptides that differ in one or few amino acid residues and by the isoform-specific QconCAT peptides. SHMT isoforms were analyzed by the label-free approach only.

### The GCS concentration in *Synechocystis* cells could allow formation of a GDC

To our knowledge, GCS protein contents were only determined for pea leaf mitochondria so far (for example, Oliver *et al.*, 1990). During this study, data for heterotrophically grown Arabidopsis cells were published (Fuchs *et al.*, 2019). It was therefore important to likewise assess the abundance of GCS proteins and their relative molar ratios in a non-plant organism. To this end, we identified and quantified all proteins involved in the GCS and the pyruvate dehydrogenase complex (PDC; Table 1) in *Synechocystis* Hi3 LC-MS datasets generated earlier in our laboratory (Gärtner *et al.*, 2019). The GCS proteins, unsurprisingly, are much less abundant in *Synechocystis* than they are in photorespiring mitochondria, which however implies that a considerable fraction of L-protein is unavailable for the GCS because it is bound to the PDC. This fraction was assessed on the assumption that the mean content of the two *Synechocystis* PDC E1 alpha and E1 beta subunits equals the amount of L-protein bound to prokaryotic PDC (Patel *et al.*, 2014). Hence, within the limits of the Hi3 technique, it appears that the L-, P-, T- and H-protein are present in a molar ratio of approximately 1L_2_-0.5P_2_-0.6T-2.4H in *Synechocystis* cells and collectively represent at least ∼0.051 mole %, corresponding to ∼0.07% mass % of the total protein. With a protein content of roughly 300 mg ml^-1^ for *Synechocystis* (Jahn *et al.*, 2018), this value corresponds to about 0.2 mg ml^-1^ total GCS protein, which is distinctly above the pea leaf GDC dissociation threshold of about 0.08 mg ml^-1^ determined by (Oliver *et al.*, 1990).

**Table 1.** Quantification of GCS proteins in *Synechocystis* by quantitative proteomics. Values show relative amounts and GCS component ratios as determined by using the Hi3 technique. Means were calculated form our previous dataset (Gärtner *et al.*, 2019), which contains reliable data for all GCS and PDC subunits. The fraction of L-protein that is available for the GCS was calculated on the assumption that the molar amount of L-protein bound to prokaryotic PDC equals the average of the two PDC E1 subunits (Patel *et al.*, 2014).

|  | Mole % | Molar ratio<br>to GCS L-protein | Mass % |
| --- | --- | --- | --- |
| PDC E1alpha | 0.058 | 3.4 | 0.068 |
| PDC E1beta | 0.063 | 3.7 | 0.069 |
| PDC E2 | 0.118 | 6.9 | 0.163 |
| Total L-protein (PDC + GCS) | 0.077 | 4.5 | 0.121 |
| PDC L-protein (E3 ~ mean of E1) | 0.060 | 3.5 | 0.094 |
| **GCS L-protein** | **0.017** | **1.0** | **0.026** |
| **GCS P-protein** | **0.009** | **0.5** | **0.028** |
| **GCS T-protein** | **0.005** | **0.3** | **0.007** |
| **GCS H-protein** | **0.020** | **1.2** | **0.009** |
| **Sum GCS L-, H, T-, P-protein** | **0.051** | **3.0** | **0.070** |
| SHMT | 0.110 | 6.5 | 0.157 |

### Recombinant *Synechocystis* GCS proteins

In order to study interactions between *Synechocystis* GCS proteins that go beyond enzyme-substrate interactions it was necessary to produce appropriate amounts of pure recombinant proteins. We have earlier reported the production and properties of recombinant *Synechocystis* P- and H-protein in *E. coli* (Hasse *et al.*, 2007, Hasse *et al.*, 2013). For our present study, we additionally developed overexpression protocols for the T- and the L-protein. The overexpressed proteins were first purified by IMAC and then further purified and quality checked by size-exclusion chromatography (SEC) and SDS-PAGE, respectively (Figure 4). During SEC in the low-ion-strength GCS buffer, the P-protein eluted at about the calculated dimeric size of ∼200 kDa (Figure 4), whereas the dimeric L-protein showed an somewhat higher than predicted apparent size of ∼150 kDa. The H-protein eluted in this buffer as a multimer with an apparent size of ∼75-80 kDa, possibly a tetramer. This pattern changed in a buffer containing 50 mM NaCl, in which the P- and L-protein eluted somewhat earlier and the H-protein tetramers dissociated to form dimers. Full lipoylation of the H-protein was confirmed by using native PAGE, which separates the lipoylated holoprotein from the apoprotein. Similar to the native T-protein from other sources (Cohen-Addad *et al.*, 1997, Guilhaudis *et al.*, 2000), the *Synechocystis* T-protein forms insoluble aggregates within a few hours after purification when overexpressed alone but remains in solution when co-expressed together with H-protein. The formed complex eluted from the SEC column with an apparent size of ∼100 kDa in the GCS buffer, possibly corresponding to a TH_4_ complex. The interaction between the two proteins was weaker in the high-salt buffer, in which a smaller, likely TH_2_ complex eluted at ∼75 kDa followed by the H-protein dimer at ∼35 kDa.

**Figure 4.**
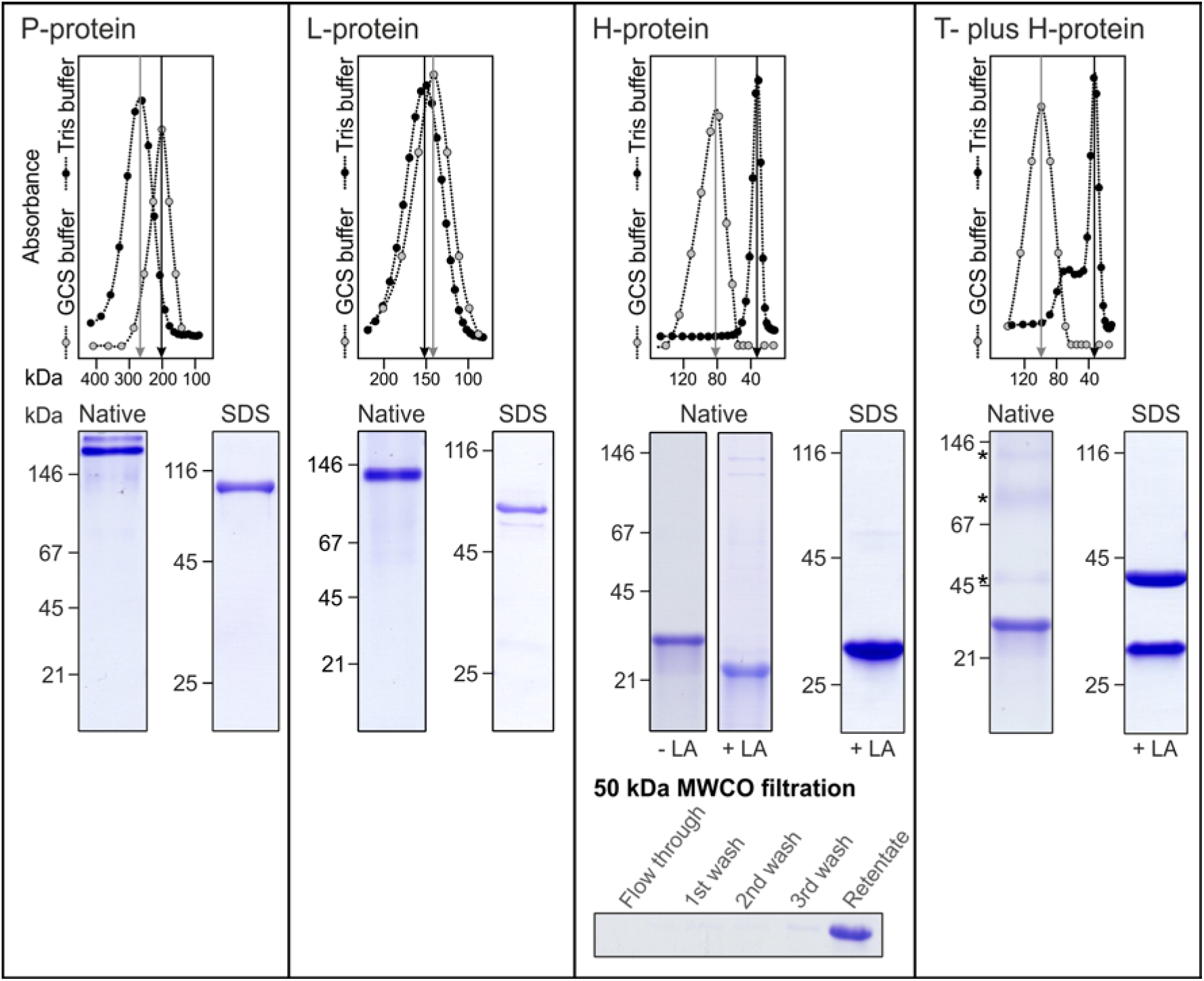
SEC purification of recombinant *Synechocystis* GCS proteins. In the low-ion-strength GCS buffer (open circles), which facilitates protein-protein interaction, the P-, L- and H-protein eluted in SEC as dimers (P_2_, L_2_) and likely tetramers (H_4_). The *Synechocystis* T-protein requires H-protein to remain in solution and elutes in the GCS buffer as an ∼100 kDa complex, likely TH_4_. The H-protein multimers and the TH_4_ complex disaggregate in a buffer containing 50 mM NaCl (closed circles). Similarly, the H-protein dissociates from the TH_4_ complex during SEC in the high-salt buffer and T-protein multimers become visible following non-denaturing PAGE (faint bands labeled with * at 45 kDa and higher). The bottom panel confirms that T- and H-protein associate in a complex larger 50 kDa, preventing passage through the 50 kDa MWCO filter membrane.

Specific maximum activities were 0.37 ± 0.01 µmol min^-1^ mg^-1^ for the P-protein (bicarbonate-exchange reaction) and 14 µmol min^-1^ mg^-1^ (with lipoic acid, K_m_ = 830 µM) or 3 µmol min^-1^ mg^-1^ (with reduced H-protein, K_m_ = 7 µM) for the L-protein. By contrast, the recombinant *Synechocystis* T-protein showed no activity, neither individually nor in the total GCS activity assay (no NADH generation, no NH_3_ release).

### Pull-down studies show interaction between the P- and L-protein

IMAC purification of the *Synechocystis* H-protein reproducibly recovered small amounts of several proteins of the overexpression host, including the *E. coli* GCS P-protein (Figure 5A), which corresponds to earlier reports that chicken and plant P- and H-protein associate to form a relatively stable P_2_H_2_ (enzyme-substrate) complex (Hiraga and Kikuchi, 1980, Walker and Oliver, 1986). By contrast, neither the T-protein nor the L-protein co-purified with H-protein. In order to test whether protein-protein interactions also occur between GCS enzyme proteins, that is, beyond enzyme-substrate interactions, we used the recombinant P-protein and L-protein as baits to recover interacting proteins from a *Synechocystis* cell lysate in pull-down experiments. The results shown in Figure 5B and C demonstrate that the immobilized *Synechocystis* P-protein specifically binds to the *Synechocystis* L-protein and vice versa. Mass spectrometry showed that several other *E. coli* and *Synechocystis* proteins were also present in the IMAC eluates of this reciprocal experiment including variable amounts of the *Synechocystis* H-protein. *Synechocystis* T-protein was not identified in either eluate, possibly because this particular GCS protein is mostly bound to the membranes and therefore not present in the soluble *Synechocystis* protein fraction.

**Figure 5.**
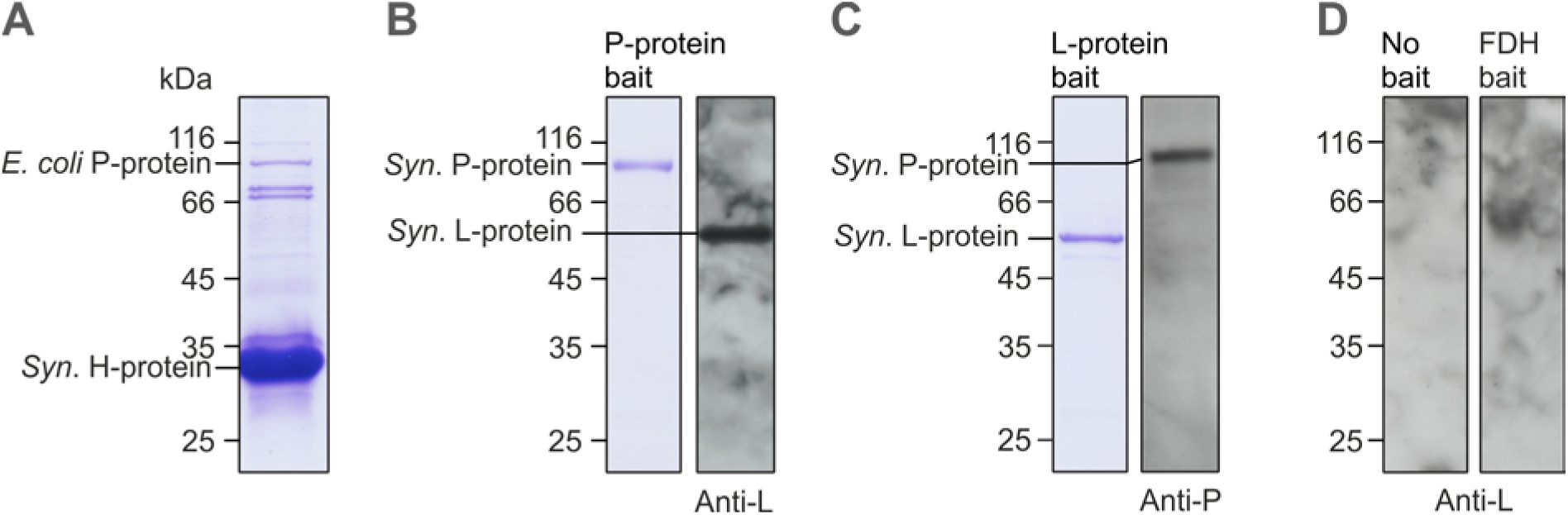
Pull-down of proteins interacting with *Synechocystis* P- and L-protein. **(A)** *E. coli* P-protein copurifies with the recombinant *Synechocystis* H-protein. **(B)** Immobilized *Synechocystis* P-protein recovers the L-protein from *Synechocystis* lysate proteins as shown by SDS-PAGE and immunoblot with a monospecific antibody. **(C)** Immobilized *Synechocystis* L-protein recovers the P-protein from *Synechocystis* lysate proteins as shown by SDS-PAGE and immunoblot with a monospecific antibody. **(D)** L-protein is not recovered from the *Synechocystis* lysate proteins in the absence of immobilized P-protein (left) or by *Pseudomonas* formate dehydrogenase as an unrelated bait (right).

### *Synechocystis* GCS proteins form a multiprotein complex at high concentration *in vitro*

The pea leaf GDC, which is the only GDC investigated so far, disaggregates at low and reassembles at high protein concentration (Neuburger *et al.*, 1986, Oliver *et al.*, 1990). We therefore examined whether the *Synechocystis* GCS proteins are likewise able to form a multiprotein complex at an appropriate concentration. As expected, all four individual GCS proteins with the 214 kDa P-protein dimer as the largest protein completely permeated a size-selective 300 kDa MWCO filter membrane (Figure 6A). We next mixed the four GCS proteins at a combined concentration above the level known to trigger formation of the plant GDC. Under this condition, substantial fractions (approximately 36% in total) of all four proteins were retained on the filter membrane even after three successive washes with fresh buffer, demonstrating formation of a multiprotein complex comprising all four GCS proteins (Figure 6B and C). The protein composition of the retentate did not noticeably change when we added more H-protein or used the (non-lipoylated) H-apoprotein (not shown). This confirms that H-protein was not limiting in our experiments and suggests the lipoyl arm is not essential for complex formation, corresponding to the formation of rather stable H-protein tetramers as shown in Figure 4. The cyanobacterial ‘GDC’ forms rapidly, even without preincubation before MWCO filtration (Figure 6B), but complex formation requires distinctly more than 10 minutes to approach equilibrium (Figure 6D). Whilst complex formation has an absolute requirement for T-protein (Figure 6E), L-protein is not necessary for the formation of a less stable P-T-H complex larger than 300 kDa (Figure 6F). Uncalibrated densitometric scans from seven SDS-PAGE patterns (independent experiments with different protein preparations) indicate an approximate molar GCS protein ratio in the retentate of 1L_2_-0.25P_2_-3T-26H (Figure 6G).

**Figure 6.**
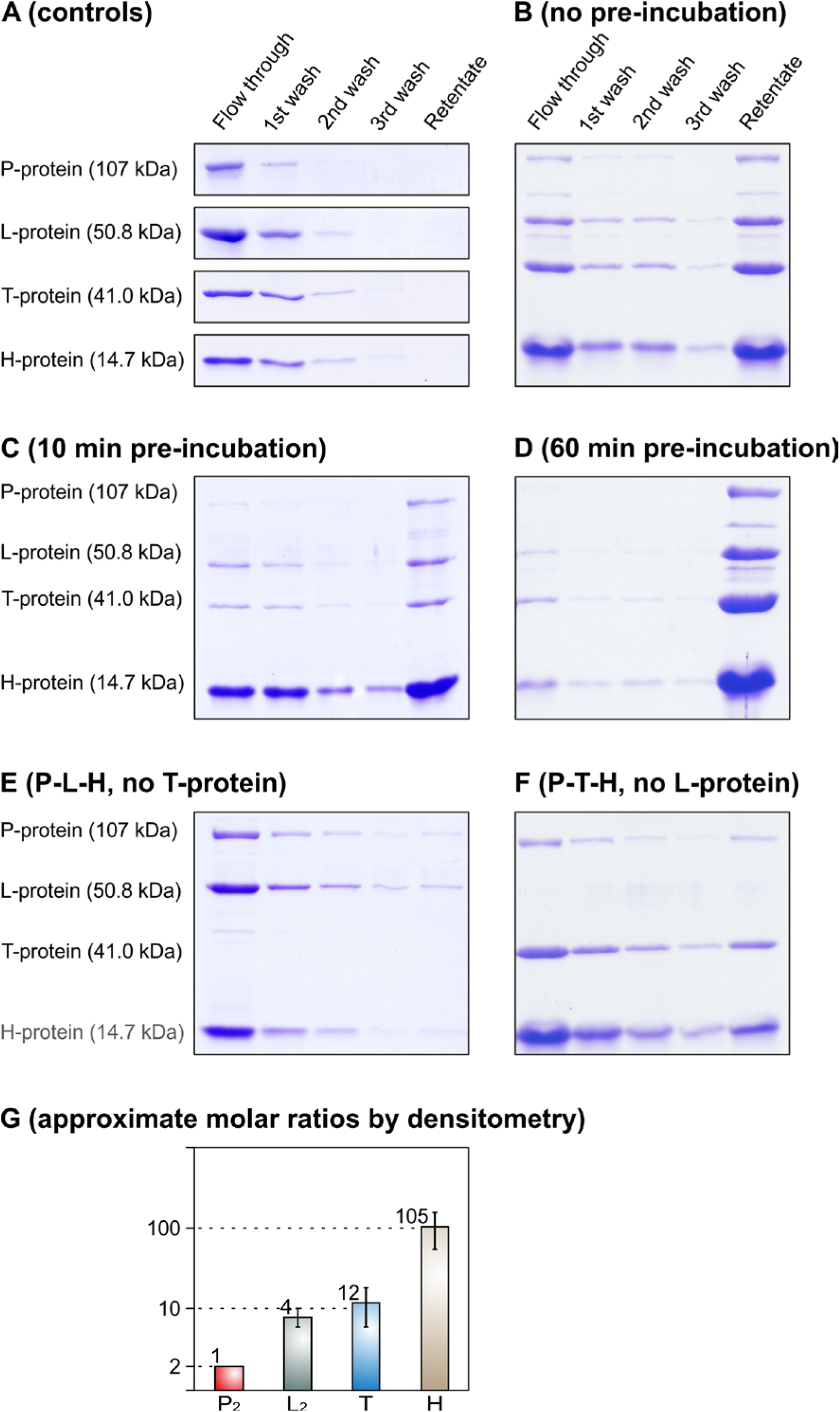
*Synechocystis* GCS proteins form a complex larger than 300 kDa *in vitro*. **(A)** All GCS proteins permeate a 300 kDa MWCO filter membrane if individually applied. Next to the first filtration, the retentate was three times rediluted with 50 µl GCS/Triton buffer and re-filtrated. None of the GCS proteins can be detected in the final retentate after three successive washes as examined by SDS-PAGE. Initial concentrations were 5.5 µg P-protein, 45 µg H-protein, 14 µg T-protein or 11 µg L-protein in 50 µl GCS/Triton buffer. **(B)** If applied as a mixture preincubated for 10 min under otherwise identical conditions, substantial fractions of all four GCS proteins are retained on the 300 kDa MWCO filter membrane. The initial combined protein concentration was 1.5 mg ml^-1^. **(C)** The artificial P-H-T-L complex forms rapidly even without preincubation. Other conditions as in B. **(D)** P-H-T-L complex formation is more complete after 60 min preincubation. Other conditions as in B. **(E)** P-H-T-L complex formation requires T-protein. Other conditions as in B. **(F)** A P-T-H complex larger then 300 kDa forms without L-protein but is less stable than the P-H-T-L complex. Other conditions as in B. **(G)** Approximate molar ratios (4L_2_-1P_2_-12T-105H) for the artificial *Synechocystis* GDC as calculated from SDS-PAGE (non-calibrated densitometry, P-protein arbitrarily set to 2 (one P_2_ dimer), seven independent repeats of the experiment shown in panel B).

## DISCUSSION

### New and variable stoichiometry of the plant GDC

It was early suggested that the four GCS proteins might interact with each other, beyond simple enzyme-substrate interactions, to form a rather unstable complex *in vitro* (for example, Hiraga *et al.*, 1972) but it was not known whether this feature has any physiological significance. Later, such a GDC could be isolated from the matrix of pea leaf mitochondria by cautious disruption followed by 300 kDa MWCO filtration, demonstrating for plant leaves that nearly all GCS proteins are bound in a stable complex *in organello* (Neuburger *et al.*, 1986, Oliver *et al.*, 1990). This breakthrough was possible because pea leaf mitochondria contain very large amounts of GCS proteins (∼30% w/w, ∼130 mg ml^-1^; Oliver *et al.*, 1990, Vauclare *et al.*, 1996). This previous estimate is close to the GCS concentration of approximately ∼44% (w/w) in the Arabidopsis mitochondrial matrix that we determined from Hi3 data (Supplementary Table S 1). At a crude total matrix protein concentration of 0.25 mg ml^-1^ in vitro, corresponding to about 0.08 mg ml^-1^ (2.6 µM) total GCS proteins, the GDC dissociates into the four individual component proteins (Oliver *et al.*, 1990).

ELISA data from the same group suggested an approximate component ratio of the pea leaf GDC of 1L_2_-2P_2_-9T-27H (Oliver *et al.*, 1990). This is the only GDC stoichiometry reported so far, which prompted us to re-examine molar ratios for the pea GDC protein components in comparison with the Arabidopsis GDC by mass spectrometry. We did so by using three different approaches: label-free Hi3 quantification (Silva *et al.*, 2006), the QconCAT technology (Pratt *et al.*, 2006), and quantification by isotope-labeled proteotypic peptides (SpikeTides_TQL, Schnatbaum *et al.*, 2011). The QconCAT technology produced the most comprehensive data with calculated molar ratios of about 1L_2_-4P_2_-8T-26H and 1L_2_-4P_2_-8T-20H for pea and Arabidopsis, respectively, which except the twofold higher P-protein content in our data is close to the ELISA results reported by Oliver et al. (1990) mentioned above. The minimum mass of a multiprotein GDC of this composition would be 1,550-1,650 kDa. The different H-protein contents are remarkable insofar as they suggest the GDC’s stoichiometry may vary between species; however, such hypothesis would assume all GCS proteins are GDC-bound in photorespiring mitochondria. It is interesting to note that the above stoichiometry likewise does not very well correspond to a recent study, which suggests a 4P_2_-26T-25H ratio in heterotrophically grown Arabidopsis cells (Fuchs *et al.*, 2019). A possible reason for the difference between plant species and photorespiring versus heterotrophic cells could be coexistence of complexed and freely diffusing GCS proteins, particularly at concentrations close to the GDC association/dissociation threshold.

Data obtained by using the other two techniques corroborated this stoichiometry, despite the specific methodical difficulties that P-protein could not be quantified by using the isotope-labeled synthetic peptide and H-protein was underestimated by the label-free Hi3 quantification. The latter effect was not unexpected because of the small number of H-protein tryptic peptides (Supplementary Figure S 3) that can be used to calculate the average abundance of the three most intense peptide signals in the Hi3 approach. It shall be noted that the GDC L-protein was slightly overestimated in Oliver *et al.* (1990) and our present experiments, because L-protein is also a component of 2-oxoacid decarboxylase complexes, such as the PDC (Bourguignon *et al.*, 1996). PDC however binds only about 12% of total mitochondrial L-protein as calculated from the Hi3 shotgun proteomics data (Supplementary Table S 1, assuming one PDC-E3 per five PDC-E1 subunits). Therefore, a correction of the above plant GDC stoichiometries for the mitochondrial PDC L-protein is not needed at this stage and additionally would be difficult because the subunit stoichiometry of plant leaf mtPDC is not yet exactly known (Mooney *et al.*, 2002, Patel *et al.*, 2014).

We next determined relative contributions of the different protein isoforms to the Arabidopsis GDC and the cooperating enzyme SHMT. These results were highly consistent in comparison of the QconCAT-based and label-free approaches (Figure 3). P-protein and L-protein are mostly, each to about 80%, represented by the isoforms GCS-P1 (At4g33010) and mtLPD1 (At1g48030). In case of the L-protein, this supports earlier studies using other techniques (Luethy *et al.*, 2001, Lutziger and Oliver, 2001). The H-protein isoforms GCS-H1 (At2g35370) and GCS-H3 (At1g32470) are present in about equal amounts, suggesting they are functionally redundant. GCS-H2 (At2g35120) was not detected in leaf mitochondria. This protein is exceptional because it shares only limited (about 60%) identity with GCS-H1 and GCS-H3 and is the only H-protein expressed in Arabidopsis roots and essential for seed development and maybe other processes (Bauwe, 2018, and unpublished data). SHMT1 (At4g37930) could well be the only SHMT in mesophyll mitochondria because the second mitochondrial isoform, SHMT2 (At5g26780) cannot be imported (Engel *et al.*, 2011). The very small fraction of this isoform hence indicates that leaf mitochondria preparations are mostly mesophyll mitochondria although they inevitably also contain small amounts of mitochondria originating from other leaf tissues, such as the vascular bundle and others.

### Is there a GDC in cyanobacteria?

By contrast to the very high concentration of GCS proteins in leaf mitochondria, heterotrophic plant cells and organs, and cyanobacteria contain much less (for example, Kopriva *et al.*, 1995, Fuchs *et al.*, 2019). It is not known whether the GCS operates in a structurally organised mode in such mitochondria and prokaryotic cells as it does in photorespiring mitochondria or whether the GCS enzymes diffuse freely, being kinetically linked by their shared mobile H-protein substrate.

In order to test this, we chose the *Synechocystis* GCS, which is considered a *bona fide* cyanobacterial model for the eukaryotic GCS (Hasse *et al.*, 2013). Since the plant GDC begins to disaggregate below 0.08 mg ml^-1^ total GCS protein (0.25 mg ml^-1^ crude matrix protein; Oliver *et al.*, 1990), corresponding to a combined total molar concentration of about 2.6 µM GCS proteins, we checked whether the GCS proteins are present in a similar or higher level in *Synechocystis* cells and what their molar ratios are. To this end, we re-examined a Hi3 dataset available in our laboratory from earlier experiments (Gärtner *et al.*, 2019), which contained quantitative data for all GCS proteins and PDC subunits. From these previously collected data, we calculated an approximate molar ratio of 1L_2_-0.5P_2_-0.6T-2.4H in *Synechocystis* cells (Table 1); however, there is yet no independent support for this ratio from other methods such as QconCAT or spiked peptide quantification. This is critical only for the *Synechocystis* H-protein, which likely is largely underestimated by Hi3 quantification due to the inherent difficulties in the quantification of small proteins, represented by very few tryptic peptides. This can be seen from the fact that a similar ratio of approximately 1L_2_-0.2P_2_-0.4T (H-protein could not be quantified) can be calculated from the independent *Synechocystis* proteome data of Zavřel *et al.* (2019).

We also calculated from the proteome data of Gärtner *et al.* (2019) that the GCS proteins would collectively represent approximately 0.07% (w/w) of the total protein in *Synechocystis* (Table 1) and a larger fraction in the soluble protein fraction. Similar levels can be derived from proteome data of *Synechocystis* provided by Jahn *et al.* (2018) and Zavřel *et al.* (2019). With a mean protein content of about 300 mg ml^-1^ for *Synechocystis* (Jahn *et al.*, 2018), this fraction corresponds to a combined concentration of the GCS proteins of about 0.2 mg ml^-1^, which is surprisingly close to and even somewhat less than the recently reported GCS concentration in heterotrophic Arabidopsis cells (Fuchs *et al.*, 2019) and 2.5-fold higher than the above mentioned dissociation threshold of 0.08 mg ml^-1^ for the pea leaf GDC (Oliver *et al.*, 1990). It hence appears that a GDC could form in *Synechocystis* (and heterotrophic plant) cells.

In order to test this hypothesis further, we overexpressed and purified His-tagged versions of the four *Synechocystis* GCS proteins. These proteins were examined for individual multimerisation and for pairwise and multiple interactions, particularly their potential to form large complexes. For the *Synechocystis* H-protein, SEC, and retention by a 50 kDa MWCO filter (Figure 4) confirmed our earlier finding (Hasse *et al.*, 2007) that this protein forms relatively stable multimers (dimers in high salt and tetramers in low salt). Formation of H-protein tetramers is not specific for *Synechocystis* but was also observed with the H-protein from *Peptococcus glycinophilus* (Robinson *et al.*, 1973) and to some extent with plant H-protein (Oliver *et al.*, 1990).

Given its function as a shared substrate, it is unsurprising that the H-protein binds to each of the three GCS enzyme proteins of *Synechocystis* (Figure 4, T-H_2_ or T-H_4_; Figure 5A, P-H; Figure 6F, P-T-H). The stability of these associations is remarkable; some of them had been observed with the GCS from other sources. For example, two monomers of chicken liver H-protein bind fairly stably per one P-protein dimer (Hiraga and Kikuchi, 1980, Kikuchi and Hiraga, 1982), whereas H- and T-protein monomers from chicken liver (Okamura-Ikeda *et al.*, 1982, Okamura-Ikeda *et al.*, 2010) and pea (Cohen-Addad *et al.*, 1997) form stable T_1_H_1_ complexes, the latter of which has been crystallized (Guilhaudis *et al.*, 2000). Direct interaction between the plant H-protein and L-protein, except via the freely exposed reduced lipoyl arm during catalysis, was not observed (Faure *et al.*, 2000, Neuburger *et al.*, 2000). That said, due to the binding of H-protein monomers to the P_2_- and the T-protein, formation of the full four-protein plant GDC would require direct physical interaction between the P-, the T-, and the L-protein or, less likely, of these proteins with a hypothetical core 30mer of H-protein as suggested by Oliver and Raman (1995). Indeed, liver mitochondria P- and L-protein have been shown to bind tightly to one another without any apparent involvement of H-protein (Hiraga *et al.*, 1972, Motokawa and Kikuchi, 1972). These findings with eukaryotic GCS proteins correspond well to our observation that the *Synechocystis* P- and L-protein are mutual interaction partners (Figure 5B, C). However, we presently do not exclude that the P_2_-L_2_ interaction at least partly could include or even be based on multi-way-binding H-protein tetramers, the presence of which seems to be a characteristic feature of the *Synechocystis* GCS.

Following Neuburger et al. (1986), we considered retention on a 300 kDa MWCO filter as evidence for the formation of a *Synechocystis* GCS multiprotein complex. Indeed, by contrast to the easily permeating individual proteins, an L-P-T-H complex larger than 300 kDa formed rapidly when the four proteins were present as a mix at a combined concentration of 1.5 mg ml^-1^. Under this condition, the filter retained about one third of the applied protein after several washes, demonstrating formation of a multiprotein complex comprising all four GCS proteins. When the L-protein was omitted, the other three GCS proteins formed a less stable complex larger than 300 kDa; however, presence of T-protein is essential for complex formation. The stoichiometry of the *Synechocystis* GDC formed *in vitro*, if it is fixed at all, could roughly match with the 1L_2_-0.25P_2_-3T-26H ratio from densitometric analysis. The difference between this ‘stoichiometry’ and the whole cell GCS protein ratio of 1L_2_-0.5P_2_-0.6T-2.4H, particularly with respect to the T- and maybe the H-protein, could indicate coexistence of free and complex-bound GCS proteins *in vivo* as discussed above as a possibility for heterotrophically cultured Arabidopsis cells. Additionally, in comparison with the plant GDC molar protein ratios of about 1L_2_-4P_2_-8T-20/26H, it is well possible that the composition of a (still hypothetical) *Synechocystis* GDC could differ from that of the plant leaf GDC.

The recombinant *Synechocystis* T-protein is soluble only in the presence of H-protein and therefore difficult to handle, a feature by which it resembles the native T-protein from other sources (Cohen-Addad *et al.*, 1997, Guilhaudis *et al.*, 2000, Okamura-Ikeda *et al.*, 2010). This has been interpreted as a possible chaperone function of the H-protein in protecting the T-protein from inactivation (Cohen-Addad *et al.*, 1997). Unfortunately, the recombinant *Synechocystis* T-protein did not show enzymatic activity. The reason for this is unclear given that repeated sequencing had confirmed correctness of the respective nucleotide sequence in the expression vector. The T-protein’s N-terminus is essential for proper binding of H-protein (Lee *et al.*, 2004), which may be blocked by the N-terminal His-tag in the recombinant T-protein. This is not very likely, however, because our experiments, such as the overexpression and SEC (Figure 4), confirm binding of the H-to the T-protein to form a soluble T-H_2_ or T-H_4_ complex. Alternatively, competitive binding of excess oxidised H-protein (the H-protein is produced in *E. coli* in the dithiolane form (Macherel *et al.*, 1996)) could prevent association with the aminomethylated form of the H-protein (Guilhaudis *et al.*, 2000), inhibiting the T-proteins enzymatic activity. Whatever the reason is, solving this methodical hurdle will be essential for the further examination of the effects that the formation of a GDC has on GCS activity in the *Synechocystis* system.

## CONCLUSIONS

By the application of three quantitative mass spectrometry techniques, we calculated molar ratios of the leaf mesophyll GDC component proteins of 1L_2_-4P_2_-8T-26H and 1L_2_-4P_2_-8T-20H for pea and Arabidopsis, respectively, with a less than 20% overestimation of L_2_ available for the GDC. Our new data indicate a twofold higher P-protein content of the GDC than the ELISA-based stoichiometry reported by Oliver et al. (1990). The minimum mass of a multiprotein GDC of this composition would be 1,550-1,650 kDa. In Arabidopsis leaves, the GDC contains four times more of the isoproteins GCS-P1 and GCS-L1 relative to GCS-P2 and GCS-L2, respectively, whereas the H-proteins GCS-H1 and GCS-H3 are present in about equal amounts, suggesting they are functionally redundant. GCS-H2 is not a component of the leaf mesophyll GDC.

The four *Synechocystis* GCS proteins associate *in vitro* to form a multiprotein complex larger than 300 kDa. The GCS content of *Synechocystis* cells is about 0.2 mg protein ml^-1^, which, by analogy to the reported dissociation threshold of 0.08 mg ml^-1^ of the plant GDC, could trigger complex formation *in vivo*. The determined L-P-T-H ratios in *Synechocystis* suggest that such complex would be different from the leaf mesophyll GDC and likely coexist with mobile GCS proteins. We also speculate that the (still hypothetical) *Synechocystis* GDC could involve H-protein multimers to crosslink the three GCS enzyme proteins, resulting in multienzyme metabolic complexes that facilitate glycine metabolism.

## MATERIAL AND METHODS

### Mitochondria

Mitochondria were isolated from leaves of *Pisum sativum* cv. Kleine Rheinländerin (3-weeks-old plants) and *Arabidopsis thaliana* Col-0 (8-weeks-old plants) by Percoll density gradient centrifugation according to Keech et al. (2005). Mitochondria were stored at −80 °C for enzyme measurements in a buffer containing 300 mM sucrose, 10 mM TES, 10 mM K-phosphate, 2 mM Na-EDTA, pH (KOH) 7.5, and for mass spectrometry in a buffer containing 50 mM Tris-HCl, pH 7.5. Matrix proteins were released by several freeze-thaw cycles followed by 1 h centrifugation at 40 000 g, 4 °C according to Neuburger et al. (1986). Purity was confirmed and the GCS content quantified by label-free absolute quantification of proteins by the Hi3 method as described below.

### Synechocystis

*Synechocystis* sp. PCC 6803 cells were grown in BG11 medium (Rippka *et al.*, 1979) at 29°C, 5% CO_2_, 120 µE m^2^ s^-1^. Cells were lysed and proteins were extracted and quantified as described earlier (Gärtner *et al.*, 2019).

### Mass Spectrometry

Mass spectrometry methods are detailed in Supplementary Methods 1. In short, the QconCAT was generated as described in detail in Pratt et al. (2006). Following the selection of suitable proteotypic peptides (Supplementary Figure S 1), the QconCAT encoding gene was synthesized by a company (BaseClear, Leiden, NL), inserted into the expression vector pET28a and expressed in *E. coli*. The isotope-labeled form was obtained by growing cells with ^15^NH_4_Cl as sole nitrogen source and purified by immobilized metal ion affinity chromatography (IMAC). Synthetic labeled peptides (SpikeTides TQL, labeled with ^13^C^15^N Lys) were purchased from JPT Peptide Technologies (Berlin, Germany). In-solution digestion of proteins was done by filter-aided sample preparation (FASP, Wiśniewski *et al.*, 2009).

LC-MS^E^ analyses were carried out using a nanoAcquity UPLC system (Waters, Manchester, UK) coupled to a Waters Synapt G2-S mass spectrometer. Peptides were separated on an analytical column (ACQUITY UPLC HSS T3, 1.8 µm, 75 µm x 250 mm, Waters) at a flow rate of 300 nl min^-1^ using a gradient from 3% to 35% acetonitrile in 0.1% formic acid over 90 min. The Synapt G2-S instrument was operated in data-independent mode (Geromanos *et al.*, 2009). By executing alternate scans at low and elevated collision energy, information on precursor and fragment ions, respectively, was acquired (referred to as MS^E^).

Progenesis QI for Proteomics version 4.1 (Nonlinear Dynamics, Newcastle upon Tyne, UK) was used for raw data processing, protein identification and MS1 intensity-based quantification. Label-free absolute quantification was performed by the Hi3 method (Silva *et al.*, 2006) with Hi3 Phos B standard (Waters) as reference. For the comparison of protein isoforms, a relative quantification approach using non-conflicting (unique) peptides was applied. For label-based quantification, the quotients of light and heavy peptide ion abundances were multiplied by the amount of heavy peptides applied to the LC column. Values for absolute quantities are calculated as fmol on column.

### Recombinant *Synechocystis* GCS proteins

The generation and structure of the *Synechocystis* H-protein (apoprotein 14,580 Da, Slr0879-pET28a) and P-protein (107,322 Da, Slr0293-pBAD/HisA) overexpression constructs was described earlier (Hasse *et al.*, 2007).

The *Synechocystis* L-protein (*slr1096*, 50,832 Da) was PCR amplified with gene-specific primers (sense 5’-<u>CTCGAG</u>ATGAGTCAGGATTTT-3’, antisense 5’-<u>GAATTC</u>TTAAACCGCCCGTTT-3’) and *Pfu* polymerase. The amplificate was ligated into pGEM-T (Promega), from which the gene was excised with XhoI/EcoRI and ligated into pBAD/HisA (Invitrogen), yielding Slr1096-pBAD/HisA. The *Synechocystis* T-protein gene (*sll0171*, 41,036 Da) was PCR-amplified using wild-type DNA, gene-specific primers (sense 5’-<u>GGATCC</u>GTGGCCAATCTTTTCCCTG-3’ and antisense 5’-<u>GAATTC</u>TTAACGAGGTTTTTGCTCGG-3’) and *Taq* polymerase. After initial cloning of the PCR amplificate into pGEM-T (Promega), the coding sequence was excised with *Bam*HI/*Eco*RI and ligated downstream of the His tag into the multiple cloning site MCS1 of pETDuet-1 (ampicillin resistance; Novagen), yielding Sll0171-pETDuet-1. All construct were verified by sequencing.

For protein expression, transgenic *E. coli* LMG194 (P- and L-protein in pBAD/HisA) and *E. coli* BL21 DE3 (all other constructs) were grown in 2YT medium and induced with 1 mM isopropyl-β-D-thiogalactopyranoside (for T- and H-protein) or 0.2% arabinose (for L- and P-protein) at OD^600^ = 0.8-1.0 and further cultivated at 30 °C for 12h, except T-protein (25 °C/12 h). Expression of T-protein alone produced flocculating aggregates during IMAC purification, whereas coexpression with H-protein from Slr0879-pET28a produced stably soluble T-H complexes. 0.24 mM lipoic acid were added for individual or combined H-protein overexpression. The relevant features of the used overexpression systems and conditions are summarised in Supplementary Table S 3.

Cells were pelleted by centrifugation (5 min, 9 000 rpm, 4°C) and resuspended in ice-cold protein-specific buffers: 20 mM Tris-HCl pH 7.8, 50 mM NaCl, 10 mM imidazole for H- and L-protein, 50 mM Tris-HCl pH 7.5, 200 mM NaCl, 0.1% Tween 80, 1 mM dithiothreitol (DTT) for T-protein, and 20 mM Na-phosphate pH 7.8, 500 mM NaCl, 0.2 mM PLP, 15 mM β-mercaptoethanol for P-protein. Lysates were obtained by sonication (six 10 s bursts, 90 W, ice-cooling) and centrifugation (14 000 rpm, 40 min, 4°C) and used for IMAC purification as above. Columns were washed three times with 5 ml each of 20 mM Tris-HCl, 50 mM NaCl and 40 mM imidazole, pH 7.8 (H- and L-protein), 50 mM Tris-HCl pH 7.5, 500 mM NaCl, 0.1% Tween 80, 1 mM DTT (T-protein), or 20 mM Na-phosphate pH 7.8, 500 mM NaCl, 0.2 mM PLP, 15 mM β-mercaptoethanol, 40 mM imidazole (P-protein). The recombinant proteins were eluted with three 1 ml washes with the same buffers except higher imidazole concentrations (T-protein 150 mM, H- and L-protein 300 mM, P-protein 500 mM).

### Size-exclusion chromatography and immunoblotting

The IMAC-purified recombinant proteins were re-buffered and further purified on a HiLoad Superdex 200 16/600 column equilibrated in the low ion-strength GCS buffer containing 5 mM 3-(N-morpholino)propanesulfonic acid (MOPS), 5 mM Tris-HCl, pH 7.0, 1 mM serine and 20 µM PLP or, as indicated, in a high-salt buffer containing 20 mM Tris-HCl (pH 8.0) and 50 mM NaCl in an ÄktaPrime or ÄktaExplorer system (GE Healthcare). The column was calibrated with bovine erythrocyte carbonic anhydrase, bovine serum albumin and yeast alcohol dehydrogenase (Sigma Aldrich). 1 ml samples of the IMAC-purified proteins were separated at a flow rate of 1 ml min^-1^. Fractions of 1 ml were collected and analysed by polyacrylamide gel electrophoresis (SDS-PAGE, Laemmli, 1970) in combination with immunoblotting using mono-specific antibodies generated against *Flaveria trinervia* H-protein, *Synechocystis* T-protein, Arabidopsis L-protein and *Synechocystis* P-protein. The eluates were concentrated on Vivaspin 500 10 kDa molecular weight cut-off (MWCO) filter/concentrator columns (Sartorius) and the proteins stored at −20°C (1 to 5 mg ml^-1^, 10% glycerol).

### Molecular mass cut-off filtration

The GCS proteins were individually or in combination diluted in 50 µl GCS buffer plus 1 mM Triton X-100 (GCS/Triton buffer) and incubated for 10 min at room temperature. These samples were filtered through Vivaspin 500 50 kDa (H-protein multimerisation) or 300 kDa (all other complex formation experiments) MWCO size-selective filters/concentrators for 3 min at 15 000 g, 4 °C. The filtrates were collected and the retentates rediluted in 50 µl of the same buffer and size-filtered as before. This process was three times repeated and the three consecutive permeates and the final retentate collected. Protein concentrations were determined (Roti-Nanoquant Roth, Bradford, 1976) and all samples analyzed by SDS-PAGE on 12% polyacrylamide gels (Laemmli, 1970). The ImageJ program was used for quantitative densitometry.

### Pull-down studies

50 µl ProBond™ nickel-chelating resin (Invitrogen) was saturated in a 500 µl batch volume with 0.5 or 5 mg recombinant *Synechocystis* L-protein or P-protein, respectively, in GCS/Triton buffer or incubated with GCS/Triton buffer without bait protein as the control. After one wash cycle with the same buffer to remove unbound protein, the loaded matrices and the unloaded control matrix were incubated for 60 min with wild-type *Synechocystis* proteins obtained as a total lysate from cells collected by centrifugation from a 50 ml culture at 2-3 OD^750^, supended in GCS/Triton buffer and exposed to two french press cycles. After three washes with 500 µl each of 20 mM Tris-HCl, pH 7.8, 1 M NaCl and 40 mM imidazole, bound proteins were eluted with 150 µl of 20 mM Na-phosphate, pH 7.8, 500 mM NaCl and 300 mM imidazole, examined by SDS-PAGE in combination with immunoblotting and further analysed by mass spectrometry. ProBond™ resin saturated with recombinant formate dehydrogenase (*Pseudomonas* sp. 101) followed by incubation with wild-type *Synechocystis* lysate served as an additional control.

### Enzyme assays

P-protein activity was measured by the ^14^C-bicarbonate-exchange method according to Hasse et al. (2007). Briefly, 7.5 µg P-protein (0.08 µM) and 30 µg H-protein (2 µM) were added to a buffer containing 50 mM Na-phosphate (pH 7.0), 1 mM DTT, 0.1 mM PLP and 20 mM glycine. The reaction was initiated by adding 30 mM Na-^14^C-bicarbonate (2.5 µC) in a final volume of 900 µl at 30 °C. 270 µl samples were removed at 0, 15 and 30 min and mixed with 80 µl 50% trichloroacetic acid to stop the reaction. Control assays did not contain glycine.

L-protein activity was determined as the NAD^+^-dependent oxidation of H-protein reduced prior the assay with 70 mM tris-(2-carboxyethyl)-phosphine (TCEP) or the NADH-dependent reduction of lipoic acid. Assays contained 10 µg of L-protein in a final volume of 1 ml 100 mM K-phosphate (pH 6.3), 1.5 mM Na-EDTA, 0.6 mg ml^-1^ bovine serum albumine, 0.2 mM NAD^+^ or 0.1 mM NADH and variable substrate concentrations at 25 °C.

## ACKNOWLEDGEMENTS

We would wish to thank Drs. Shanshan Wang and Markus Wirtz (both at the University of Heidelberg) for their practical support and helpful discussions and appreciate critical discussions with Dr. Eva-Maria Brouwer (University of Rostock).

## SUPPLEMENTARY DATA

### Supplementary Methods 1. Mass spectrometry

#### Peptides for stable isotope-based quantification of plant GCS proteins and design of a QConCAT

A synthetic protein was designed that comprises a concatenation of proteotypic peptides (QconCAT, Pratt *et al.*, 2006) to facilitate the stoichiometric analysis of the GCS in Arabidopsis and pea leaf mitochondria. Peptides were selected that allowed identification of GCS proteins in previous LC-MS^E^ measurements. To achieve high comparability between Arabidopsis and pea, pairs of homologous peptides with minimal length of nine amino acids were selected from the aligned protein sequences (for example, Supplementary Figure S 3). Peptides containing methionine or cysteine were usually excluded. Methionine-containing peptides were only selected if they showed minimal oxidation in the mitochondrial extracts of Arabidopsis and pea. Two peptides per protein were incorporated into the QConCAT (Supplementary Figure S 1). For the isoforms of the Arabidopsis L-, P-, and H-proteins, one peptide each was isoform-specific while the second peptide was specific for both isoforms. The flanking sequences of the peptides were derived from the native proteins. Peptides of rabbit phosphorylase B flanked by the linker sequence ASGK (Smith *et al.*, 2016) were incorporated at both ends of the QConCAT to enable quantification of the QconCAT amount via a synthetic Hi3 Phos B standard (Waters, Manchester, UK). The QconCAT-encoding gene was synthesized (BaseClear, Leiden, NL), inserted into the expression vector pET28a and expressed in *E. coli*. The stable isotope-labeled protein was obtained by growing cells with ^15^NH_4_Cl as sole nitrogen source and purified by immobilized metal ion affinity chromatography (IMAC).

The stoichiometric analysis of the GCS was alternatively performed by using synthetic labeled peptides (SpikeTides TQL, labeled with ^13^C^15^N Lys; JPT Peptide Technologies, Berlin, Germany) as shown in the Supplementary Table S 2. The SpikeTide peptide IAILNANYMAK, which is specific for the P-protein from pea and both isoforms of the P-protein of Arabidopsis, was highly oxidized and could not be used. This finding was in contrast to the much lower ratio of oxidation products of this peptide in tryptic digests of the QConCAT and native mitochondrial P-protein.

#### In-solution digestion of proteins and tagged SpikeTide peptides

20 µg of mitochondrial matrix proteins in 20 µl solubilization buffer containing 1% sodium dodecyl sulfate (SDS), 50 mM DTT and 50 mM Tris-HCl, pH 7.6 were incubated at 95°C for 5 min, cleared by centrifugation at 12000 x g for 5 min and further processed by filter-aided sample preparation (FASP, Wisniewski *et al.*, 2009) by using Microcon YM-30-filter devices (Millipore). The processing steps for detergent removal, alkylation, buffer exchange and protein digestion comprised two initial washes with a solution of 8 M urea in 0.1 M Tris-HCl pH 8.5 (buffer UA) followed by incubation with 50 mM iodoacetamide (IAA) in UA for 20 min, two washes with UA to deplete IAA and finally three washes with 50 mM ammonium bicarbonate (buffer ABC). Then, digestion with trypsin was performed at an enzyme-to-protein ratio of 1:50 in 40 µl of 50 mM ABC at 37° C for 16 h. Peptides were collected by centrifugation and fresh trypsin solution was added onto the filter for a second digestion for 2 h. After centrifugation, the combined digests were acidified with trifluoroacetic acid (TFA, final concentration 0.25% w/v), concentrated by use of a centrifugal evaporator and diluted to a final volume of 30 µl with a solution containing 2% acetonitrile and 0.1% (w/v) formic acid in water. Peptide concentrations were measured using the Qubit protein assay (Thermo Fisher Scientific, Waltham, MA, USA).

QConCAT protein was added to mitochondrial matrix proteins before FASP and was digested in parallel to the mitochondrial proteins. Tagged SpikeTide peptides (JPT Peptide Technologies, Berlin, Germany) were solubilized in a solution consisting of 80% of 0.1 M ABC and 20% acetonitrile as recommended by the producer. The peptides were either mixed in an equimolar ratio or in a weighted ratio of 1:3:10 for the L-, T- and H-protein-specific peptides. To separate the tryptic peptides from the Qtag an equal volume of 50 mM ABC containing trypsin (enzyme to peptide ratio of 1:50) was added. Peptides were digested at 37° C for 14 h, acidified with TFA, concentrated in a centrifugal evaporator and diluted with a solution containing 2% acetonitrile and 0.1% formic acid in water. Digested mixtures of the SpikeTide peptides were added to peptide preparations of mitochondrial matrix proteins prior to LC-MS.

#### Analysis by nanoLC-MS^E^

LC-MS^E^ analyses were carried out using a nanoAcquity UPLC system (Waters, Manchester, UK) coupled to a Waters Synapt G2-S mass spectrometer via a NanoLockSpray ion source. Mobile phase A contained 0.1% formic acid in water, and mobile phase B contained 0.1% formic acid in acetonitrile. Samples containing 10 ng of peptides from digested mitochondrial matrix proteins and experiment-dependent additions of QConCAT or SpikeTide peptides supplemented with 10 fmol of Hi3 Phos B standard for protein absolute quantification (Waters) were trapped and desalted using a pre-column (nanoAcquity UPLC Symmetry C18, 5 µm, 180 µm x 20 mm, Waters) at a flow rate of 10 µl min^-1^ for 4 min with mobile phase A. Peptides were separated on an analytical column (ACQUITY UPLC HSS T3, 1.8 µm, 75 µm x 250 mm, Waters) at a flow rate of 300 nl min^-1^ using a gradient from 3% to 35% B over 90 min. The column temperature was maintained at 35 °C. The SYNAPT G2-S instrument was operated in data-independent mode (Geromanos *et al.*, 2009). By executing alternate scans at low and elevated collision energy, information on precursor and fragment ions, respectively, was acquired (referred to as MS^E^).

#### nanoLC-MS^E^ data processing, protein identification and quantification

Progenesis QI for Proteomics version 4.1 (Nonlinear Dynamics, Newcastle upon Tyne, UK) was used for raw data processing, protein identification and quantification. Alignment was performed to compensate for between-run variation in the LC separation. Peptide and protein identifications were obtained by searching against databases containing 15,789 reviewed protein sequences from *Arabidopsis thaliana* and 1903 reviewed and non-reviewed protein sequences from *Pisum sativum* (UniProt release 2018_06). The sequences of rabbit phosphorylase B (P00489) and porcine trypsin were appended into these databases. Two missing cleavage sites were allowed, oxidation of methionine residues was considered as variable modification, and carbamidomethylation of cysteines as fixed modification. Additionally, variable modifications of all amino acids by ^15^N- or ^13^C^15^N-lysine were applied for analysis of samples containing QConCAT or SpikeTide peptides. The false discovery rate was set to 4%. Peptides were required to be identified by at least three fragment ions and proteins by at least six fragment ions and two peptides. Subsequently, peptide ion data were filtered to retain only peptide ions that met the following criteria: (i) identified at least two times within the dataset, (ii) ion score greater 6.0, (iii) mass error below 10.0 ppm, (iiii) at least 6 amino acid residues in length. For label-free quantification of all proteins, the Hi3 method implemented into the Progenesis QI for Proteomics workflow was applied using the Hi3 Phos B standard (Waters) as a reference (Silva *et al.*, 2006). Hi3 peptide quantification uses the sum of the signal intensities of the three most abundant peptides of each protein, divided by the sum of the signal intensities of the three most abundant peptides of the internal standard, multiplied by the amount of standard applied to the column.

The label-free calculation of the ratios between the isoforms of the L-, P- and H-protein of Arabidopsis was not carried out by the Hi3 method, but incorporated all pairs of isoform-specific peptide ions whose sequences differed only in one or a few amino acid positions (“homologous peptides”). Such peptides should generate comparable signal intensities. From the summed up abundances of the peptide ion signals of the isoforms to be compared, their molar ratio was determined.

The QConCAT added to the mitochondrial extracts was quantified using the unlabeled phosphorylase B peptide standard (Hi3 PhosB, Waters) with which the samples were supplemented before LC-MS measurements. The QConCAT concentration was calculated as the quotient of the heavy-labeled peptide abundance (QConCAT digestion) by the light peptide abundance (Hi3 PhosB standard) times the amount of Hi3 PhosB standard. From the results for the two phosphorylase B peptides integrated into the QConCAT, the mean value was calculated.

To quantify the GDC proteins by means of isotope-labeled peptides, the quotients of light and heavy-labeled peptide ion abundances were determined. If different charge states were detected for a peptide, their abundances were summed up. It was ensured that for both light and heavy-labeled peptides the same charge states were used for the calculation. The quotients of light and heavy peptide ion abundances were then multiplied by the amount of heavy peptides applied to the LC column. Thus, values for absolute quantities are calculated as fmol on column.

**Table S 1. Arabidopsis mitochondrial matrix protein identification and Hi3 quantification of two independent mitochondrial preparations, of which each four proteolytic digestions were analyzed by mass spectrometry.**

Supplementary Table S1.xlsx

**Table S 2.**
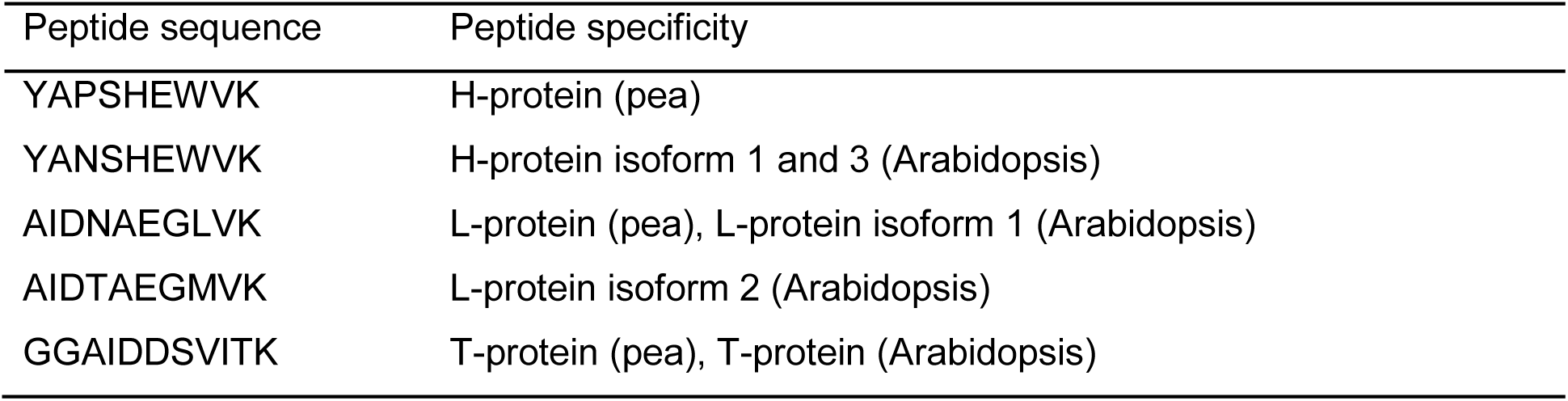
Synthetic peptides terminally labeled with ^13^C^15^N Lys (SpikeTides TQL).

**Table S 3.**
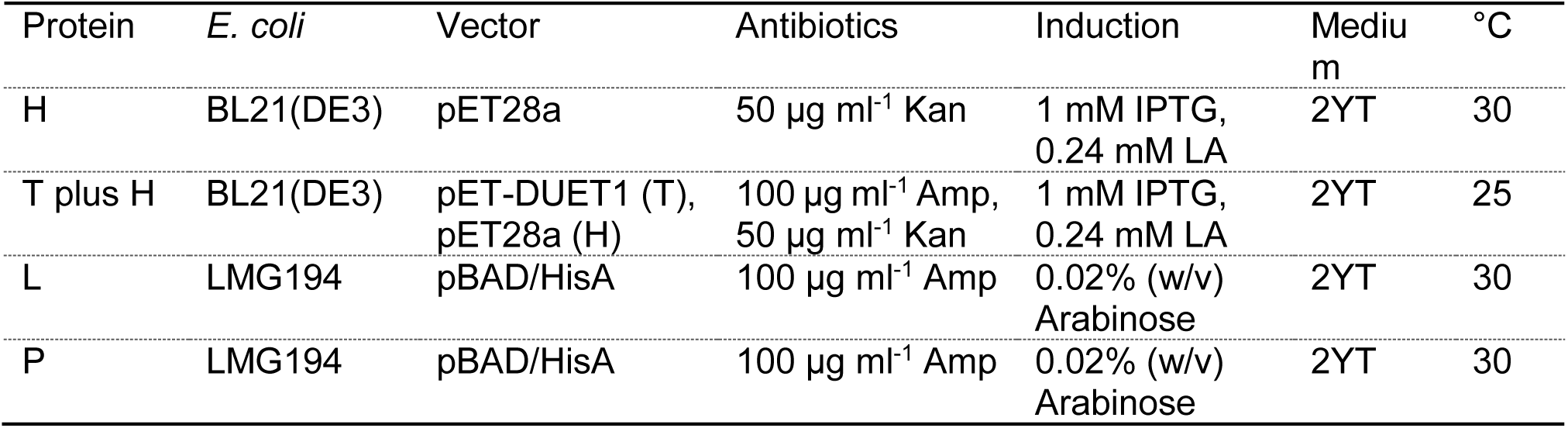
Overexpression systems. Abbreviations: Amp, Ampicillin; Kan, Kamamycin; LA, lipoic acid

**Figure S 1.**
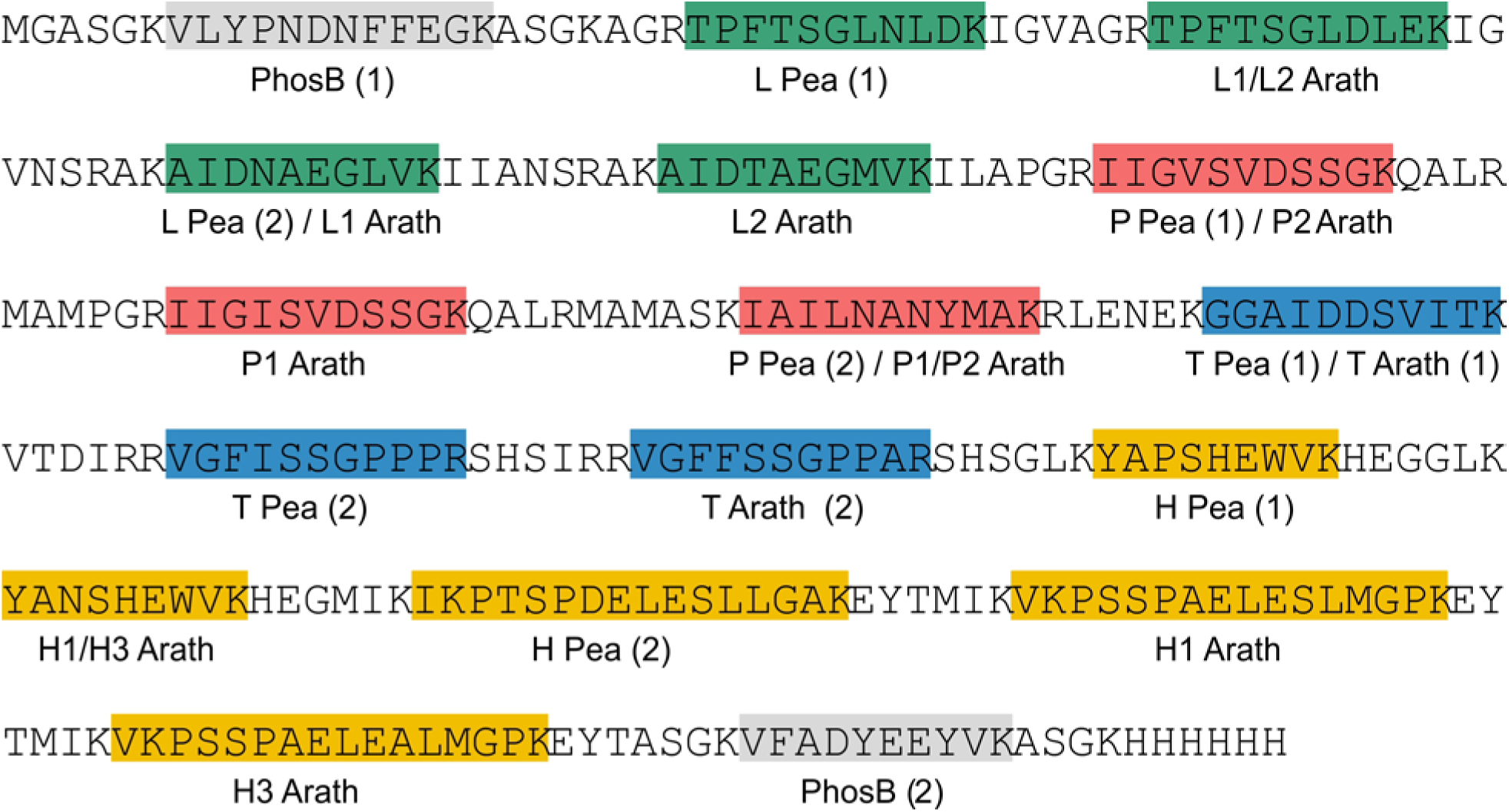
Design of the QconCAT protein. Suitable peptides were identified by analysis of LC-MS^E^ data from tryptic digests of mitochondrial matrix proteins. Two peptides per protein were selected. For the isoforms of the Arabidopsis L-, P- and H-proteins, one peptide was isoform-specific while the second peptide was specific for both isoforms. The flanking sequences of the peptides derive from the native proteins. Peptides of rabbit phosphorylase B were incorporated at both ends to enable quantification of the QconCAT amount via a synthetic Hi3 Phos B standard.

**Figure S 2.**
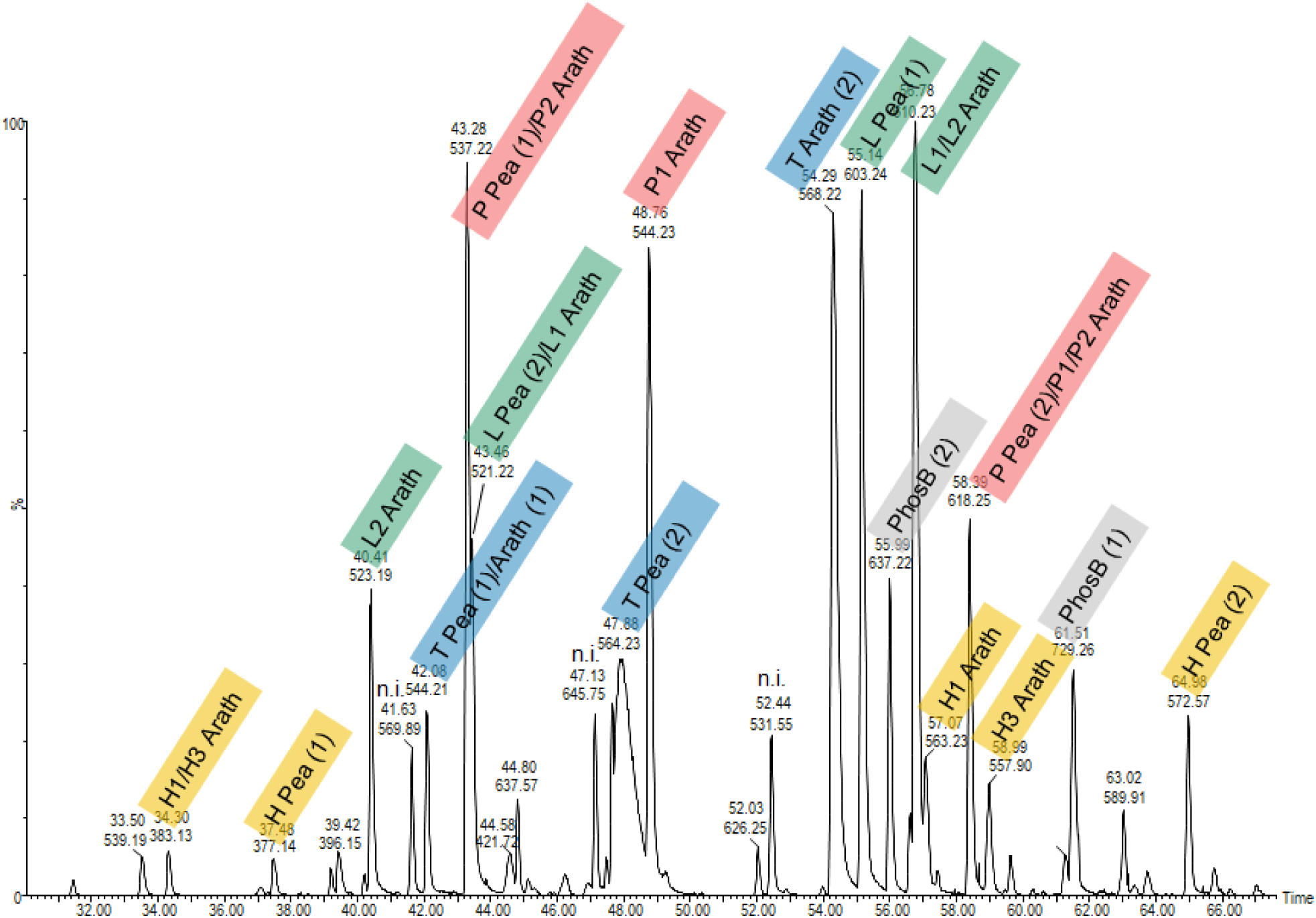
Base peak ion chromatogram of a tryptic digest of the QconCAT. The chromatogram displays all peptides included into the QconCAT.

**Figure S 3.**
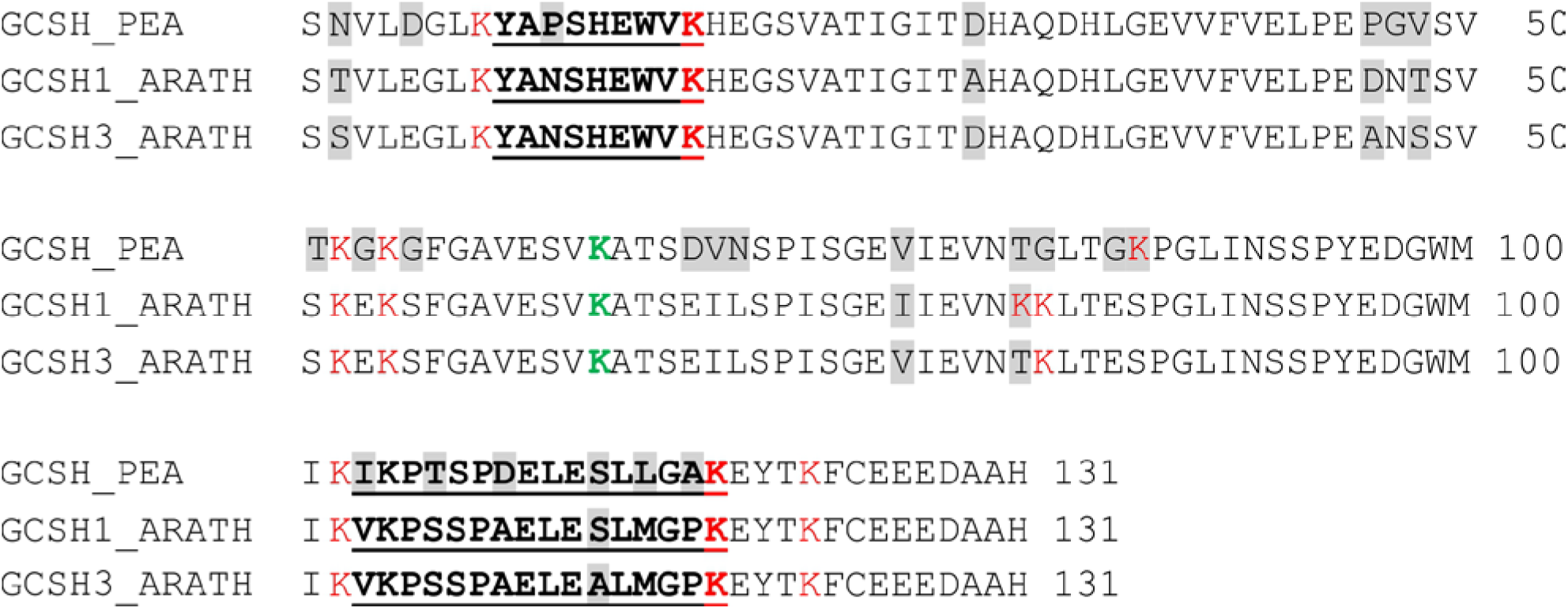
Alignment of the Arabidopsis H-protein isoforms GCS-H1 and GCS-H3 and the H-protein from pea. Amino acid sequences of the mature proteins after cleavage of the transit peptide are shown. Sequence differences are high-lighted and trypsin cleavage sites are labeled in red. The lipoyl-binding lysine is shown in green. The sequences provide only a small number of tryptic peptides, and a single detectable peptide is differentiating between the leaf mesophyll isoforms of Arabidopsis. Peptides selected for the QconCAT are underlined.

## REFERENCES

Bar-Even, A. (2016) Formate assimilation: The metabolic architecture of natural and synthetic pathways. Biochemistry, 55, 3851–3863.

Bauwe, H. (2018) Photorespiration - damage repair pathway of the Calvin–Benson cycle. In Plant Mitochondria, 2^nd^ Edition (Logan, D.C. ed). Chichester, UK: John Wiley & Sons, Ltd, pp. 293–342.

Bauwe, H., et al. (2010) Photorespiration: players, partners and origin. Trends Plant Sci., 15, 330–336.

Bauwe, H. and Kolukisaoglu, Ü. (2003) Genetic manipulation of glycine decarboxylation. J. Exp. Bot., 54, 1523–1535.

Bourguignon, J., et al. (1996) Glycine decarboxylase and pyruvate-dehydrogenase complexes share the same dihydrolipoamide dehydrogenase in pea leaf mitochondria - evidence from mass-spectrometry and primary-structure analysis. Biochem. J., 313, 229–234.

Bourguignon, J., et al. (1988) Resolution and characterization of the glycine cleavage reaction in pea leaf mitochondria. Properties of the forward reaction catalysed by glycine decarboxylase and serine hydroxymethyltransferase. Biochem. J., 255, 169–178.

Bradford, M.M. (1976) A rapid and sensitive method for the quantitation of microgram quantities of protein utilizing the principle of protein-dye binding. Anal. Biochem., 72, 248–254.

Cohen-Addad, C., et al. (1997) Structural studies of the glycine decarboxylase complex from pea leaf mitochondria. Biochimie, 79, 637–643.

Douce, R., et al. (2001) The glycine decarboxylase system: a fascinating complex. Trends Plant Sci., 6, 167–176.

Engel, N., et al. (2011) The presequence of Arabidopsis serine hydroxymethyltransferase SHM2 selectively prevents import into mesophyll mitochondria. Plant Physiol., 157, 1711-1720.

Engel, N., et al. (2007) Deletion of glycine decarboxylase in Arabidopsis is lethal under non-photorespiratory conditions. Plant Physiol., 144, 1328–1335.

Fan, J., et al. (2014) Quantitative flux analysis reveals folate-dependent NADPH production. Nature, 510, 298–302.

Faure, M., et al. (2000) Interaction between the lipoamide-containing H-protein and the lipoamide dehydrogenase (L-protein) of the glycine decarboxylase multienzyme system. 2. Crystal structures of H- and L-proteins. Eur. J. Biochem., 267, 2890–2898.

Freudenberg, W. and Andreesen, J.R. (1989) Purification and partial characterization of the glycine decarboxylase multienzyme complex from *Eubacterium acidaminophilum*. J. Bacteriol., 171, 2209–2215.

Fuchs, P., et al. (2019) Single organelle function and organization as estimated from Arabidopsis mitochondrial proteomics. Plant J., DOI10.1111/tpj.14534.

Gärtner, K., et al. (2019) Cytosine N4-methylation via M.Ssp6803II is involved in the regulation of transcription, fine-tuning of DNA replication and DNA repair in the cyanobacterium *Synechocystis* sp. PCC 6803. Front. Microbiol., 10, 1233.

Geromanos, S.J., et al. (2009) The detection, correlation, and comparison of peptide precursor and product ions from data independent LC-MS with data dependant LC-MS/MS. Proteomics, 9, 1683-1695.

Guilhaudis, L., et al. (2000) Combined structural and biochemical analysis of the H-T complex in the glycine decarboxylase cycle: evidence for a destabilization mechanism of the H-protein. Biochemistry, 39, 4259–4266.

Hasse, D., et al. (2013) Structure of the homodimeric glycine decarboxylase P-protein from *Synechocystis* sp. PCC 6803 suggests a mechanism for redox regulation. J. Biol. Chem., 288, 35333–35345.

Hasse, D., et al. (2007) Properties of recombinant glycine decarboxylase P- and H-protein subunits from the cyanobacterium *Synechocystis* sp. strain PCC 6803. FEBS Lett., 581, 1297–1301.

Hiraga, K. and Kikuchi, G. (1980) The mitochondrial glycine cleavage system. Functional association of glycine decarboxylase and aminomethyl carrier protein. J. Biol. Chem., 255, 11671–11676.

Hiraga, K., et al. (1972) Enzyme complex nature of the reversible glycine cleavage system of cock liver mitochondria. J. Biochem., 72, 1285–1289.

Jahn, M., et al. (2018) Growth of cyanobacteria is constrained by the abundance of light and carbon assimilation proteins. Cell Rep., 25, 478-486.e478.

Kawasaki, H., et al. (1966) A new reaction for glycine biosynthesis. Biochem. Biophys. Res. Commun., 23, 227–233.

Keech, O., et al. (2005) Preparation of leaf mitochondria from *Arabidopsis thaliana*. Physiol. Plant., 124, 403–409.

Kikuchi, G. and Hiraga, K. (1982) The mitochondrial glycine cleavage system. Unique features of the glycine decarboxylation. Mol. Cell. Biochem., 45, 137–149.

Kikuchi, G., et al. (2008) Glycine cleavage system: reaction mechanism, physiological significance, and hyperglycinemia. Proc. Japan Acad., Ser. B Phys. Biol. Sci., 84, 246–263.

Kisaki, T., et al. (1971) Glycine decarboxylase and serine formation in spinach leaf mitochondrial preparation with reference to photorespiration. Plant Cell Physiol., 12, 275–288.

Kopriva, S., et al. (1995) Alternative splicing results in two different transcripts for H-protein of the glycine cleavage system in the C4 species *Flaveria trinervia*. Plant J., 8, 435–441.

Laemmli, U.K. (1970) Cleavage of structural proteins during the assembly of the head of bacteriophage T4. Nature, 227, 680-685.

Lee, H.H., et al. (2004) Crystal structure of T-protein of the glycine cleavage system: Cofactor binding, insights into H-protein recognition, and molecular basis for understanding nonketotic hyperglycinemia. J. Biol. Chem., 279, 50514–50523.

Luethy, M.H., et al. (2001) Developmental expression of the mitochondrial pyruvate dehydrogenase complex in pea (*Pisum sativum*) seedlings. Physiol. Plant., 112, 559–566.

Lutziger, I. and Oliver, D.J. (2001) Characterization of two cDNAs encoding mitochondrial lipoamide dehydrogenase from *Arabidopsis*. Plant Physiol., 127, 615–623.

Macherel, D., et al. (1996) Expression, lipoylation and structure determination of recombinant pea H-protein in *Escherichia coli*. Eur. J. Biochem., 236, 27–33.

Mooney, B.P., et al. (2002) The complex fate of α-ketoacids. Annu. Rev. Plant Biol., 53, 357-375.

Motokawa, Y. and Kikuchi, G. (1971) Glycine metabolism in rat liver mitochondria. V. Intramitochondrial localization of the reversible glycine cleavage system and serine hydroxymethyltransferase. Arch. Biochem. Biophys., 146, 461–464.

Motokawa, Y. and Kikuchi, G. (1972) Isolation and partial characterization of the components of the reversible glycine cleavage system of rat liver mitochondria. J. Biochem., 72, 1281–1284.

Mouillon, J.M., et al. (1999) Glycine and serine catabolism in non-photosynthetic higher plant cells: their role in C1 metabolism. Plant J., 20, 197–205.

Neuburger, M., et al. (1986) Isolation of a large complex from the matrix of pea leaf mitochondria involved in the rapid transformation of glycine into serine. FEBS Lett., 207, 18-22.

Neuburger, M., et al. (2000) Interaction between the lipoamide-containing H-protein and the lipoamide dehydrogenase (L-protein) of the glycine decarboxylase multienzyme system. 1. Biochemical studies. Eur. J. Biochem., 267, 2882–2889.

Okamura-Ikeda, K., et al. (1982) Purification and characterization of chicken liver T-protein, a component of the glycine cleavage system. J. Biol. Chem., 257, 135-139.

Okamura-Ikeda, K., et al. (2010) Crystal structure of aminomethyltransferase in complex with dihydrolipoyl-H-protein of the glycine cleavage system: Implications for recognition of lipoyl protein substrate, disease-related mutations, and reaction mechanism. J. Biol. Chem., 285, 18684–18692.

Oliver, D.J., et al. (1990) Interaction between the component enzymes of the glycine decarboxylase multienzyme complex. Plant Physiol., 94, 833–839.

Oliver, D.J. and Raman, R. (1995) Glycine decarboxylase: protein chemistry and molecular biology of the major protein in leaf mitochondria. J. Bioenerg. Biomembr., 27, 407–414.

Patel, M.S., et al. (2014) The pyruvate dehydrogenase complexes: structure-based function and regulation. J. Biol. Chem., 289, 16615–16623.

Pratt, J.M., et al. (2006) Multiplexed absolute quantification for proteomics using concatenated signature peptides encoded by QconCAT genes. Nat. Protoc., 1, 1029.

Rippka, R., et al. (1979) Generic assignments, strain histories and properties of pure cultures of cyanobacteria. Microbiology, 111, 1–61.

Robinson, J.R., et al. (1973) Glycine metabolism. Lipoic acid as the prosthetic group in the electron transfer protein P2 from *Peptococcus glycinophilus*. J. Biol. Chem., 248, 5319–5323.

Schmitt, D.L. and An, S. (2017) Spatial organization of metabolic enzyme complexes in cells. Biochemistry, 56, 3184–3196.

Schnatbaum, K., et al. (2011) SpikeTides™ - proteotypic peptides for large-scale MS-based proteomics. Nat. Methods, 8, 272.

Silva, J.C., et al. (2006) Absolute quantification of proteins by LCMSE: A virtue of parallel MS acquisition. Mol. Cell. Proteomics, 5, 144–156.

Smith, D.G.S., et al. (2016) Design and expression of a QconCAT protein to validate Hi3 protein quantification of influenza vaccine antigens. J. Proteom., 146, 133–140.

Vauclare, P., et al. (1996) Regulation of the expression of the glycine decarboxylase complex during pea leaf development. Plant Physiol., 112, 1523–1530.

Walker, J.L. and Oliver, D.J. (1986) Glycine decarboxylase multienzyme complex. Purification and partial characterization from pea leaf mitochondria. J. Biol. Chem., 261, 2214–2221.

Wisniewski, J.R., et al. (2009) Universal sample preparation method for proteome analysis. Nat. Methods, 6, 359–362.

Yishai, O., et al. (2018) In vivo assimilation of one-carbon via a synthetic reductive glycine pathway in *Escherichia coli*. ACS Synth. Biol., 7, 2023–2028.

Zavřel, T., et al. (2019) Quantitative insights into the cyanobacterial cell economy. eLife, 8, e42508.

